# Biosynthesis of Iron Oxide Nanoparticles via *Crocus sativus* and their Antifungal Efficacy against *Verticillium* Wilt Pathogen *Verticillium dahliae*

**DOI:** 10.1101/861401

**Authors:** Tariq Alam, Fazal Akbar, Mohammad Ali, Muhammad Farooq Hussain Munis, Jafar Khan

## Abstract

Plant diseases pose threat to global food security. The excessive use of synthetic agro-chemical engender pesticide resistance. The exploration of alternative sustainable diseases management practices are crucial to overcome the devastative plant diseases. In this study, a facile innocuous approach was adopted for biogenic synthesis of Fe_2_O_3_-CsNPs via *Crocus sativus* corm aqueous extract and was evaluated for their antifungal efficacy against the *Verticillium* wilt pathogen *Verticillium dahliae*. The physico-chemical characterization of biosynthesized nanoparticles were performed through UV-visible Spectroscopy, Fourier Transform Infrared Spectroscopy, Energy Dispersive X-ray Spectroscopy, X-Ray Diffraction, and Scanning Electron Microscopy. The fungus mycelium growth was significantly inhibited in the media containing 3mg/mL Fe_2_O_3_-CsNPs. Degenerated, concentrated and shriveled hyphae were revealed in Scanning Electron Microscopy. The overall results demonstrated that the biogenic Fe_2_O_3_-CsNPs have the efficacy to control devastative phytopathogens.

## Introduction

*V. dahliae* cause *Verticillium* wilt in broad range of economic important crops. It is a polyphagous pathogen and host list is persistently expanding with diagnosis of new hosts [1-3]. Thus, the sustainable management of *Verticillium* wilt is vital. To date, the synthetic agrochemicals and pesticides has been employed for the management of wilt diseases. Nevertheless, the synthetic chemical pesticides engendered to pathogen resistance and deteriorate the ecosystem [4]. To ensure the healthy plant growth and development, sustainable diseases management practices are required. Therefore, exploration of ecofriendly and benign products are essential to overcome the pesticide resistance.

Nanoparticles possess significant antimicrobial properties and thus it has immense efficacy in the management of various diseases [5]. The antifungal and antibacterial employment of nanoparticles have been considered as economical and ecofriendly alternate strategy for the management of pathogenic microbes [6-8]. However, recently metal oxide nanoparticle are biosynthesized via environmental innocuous materials including plant extracts [9], plant tissue [10] and several other parts of the plants [11]. Iron oxide nanoparticle have attracted immense interests due their remarkable physiochemical properties. The biogenic synthesis of iron oxide nanoparticles through plants is novel approach to overcome the drawbacks of the classical methods. In the biogenic synthesis method, the phytochemicals in plants play a dual role, both as reducing and as capping agents [12, 13]. The potential of organisms to the biosynthesize nanoparticles contain both prokaryotes and eukaryotes [14]

Iron oxide nanoparticles can be exploited in broad range of application such as biomedicine, electronics, catalysis, water treatment, magnetic and environmental remediation [15-17]. Numerous plants have been employed for the biosynthesis of iron oxide nanoparticles including *Terminalia chebula*,[18] *Eucalyptus globules*,[19] Orange peels,[20] *Camellia sinensis*,[21] Sorghum beans [22] and several other plants.

Globally, *Crocus sativus* L. is the most precious medicinal plant [23]. The petal color is produced by a compound known as anthocyanins which belong to flavonoids family [24]. Many bioactive compound have been identified and isolated from saffron petals such as isorhamnetin, kaempferol, quercetin, malvidin, delphinindin and petunidin [25].

Herein, for the first time we report the biosynthesis of Fe_2_O_3_-CNPs through *Crocus sativus* corm aqueous extract and its efficacy to control *Verticillium* wilt pathogen *V. dahliae.* There is no published report on the biogenic synthesis of *V. dahliae* via *Crocus sativus* corm aqueous extract and its antiphytofungal activities. The Fe_2_O_3_-CNPs were synthesized and its antifungal efficacy against *Verticillium* wilt were evaluated. The physiochemical characterization of the synthesized nanoparticles were performed through FTIR, XRD, UV-visible, EDX and SEM.

## Materials and Methods

### 2.0.1 *Crocus sativus* aqueous extract synthesis

To prepare *Crocus sativus* corm aqueous extract, peel was thoroughly removed from the bulb, a known amount (10g) of *Crocus sativus* corm was diced into fine pieces and poured into sterile 500-mL Erlenmeyer flask. Deionized water 100 mL was poured to the pristine flask and boiled to assist in the genesis of aqueous extract. The acquired *C. sativus* aqueous extract was filtered via Whattaman No. 1 filter paper and the obtained filtrate was preserved.

### 2.0.2 Synthesis of Iron Oxide Nanoparticles

To prepare biogenic Iron Oxide Nanoparticles (Fe_2_O_3_-CsNPs), 0.1 M FeCl_3_ solution was added to the *crocus sativus* corm aqueous extract in 1:1. Fe_2_O_3_-CsNPs were subsequently acquired through the reduction method. The synthesized mixture was stirred for 60 min and then allowed to remain at room temperature for 30 minutes. The acquired colloidal suspensions were centrifuges at 10,000 rpm for 20 min and washed numerous times with ethanol, then vacuum dried at 40 °C to acquire Fe_2_O_3_-CsNPs.

### 2.0.3 Characterization of Biogenic Iron Oxide Nanoparticles

### 2.0.4 UV-visible Spectroscopic Analysis

The formation of biogenic Fe_2_O_3_-CsNPs was confirmed through UV-visible spectrometer (UV-3000 ORI, Germany). The UV-vis spectrum acquired is illustrated in Fig 1.0. The synthesized Fe_2_O_3_-CsNPs revealed the maximum absorbance peak at 318 nm. The spectrum acquired confirm the biogenic synthesis of Fe_2_O_3_-CsNPs.

**Fig 1.0.**
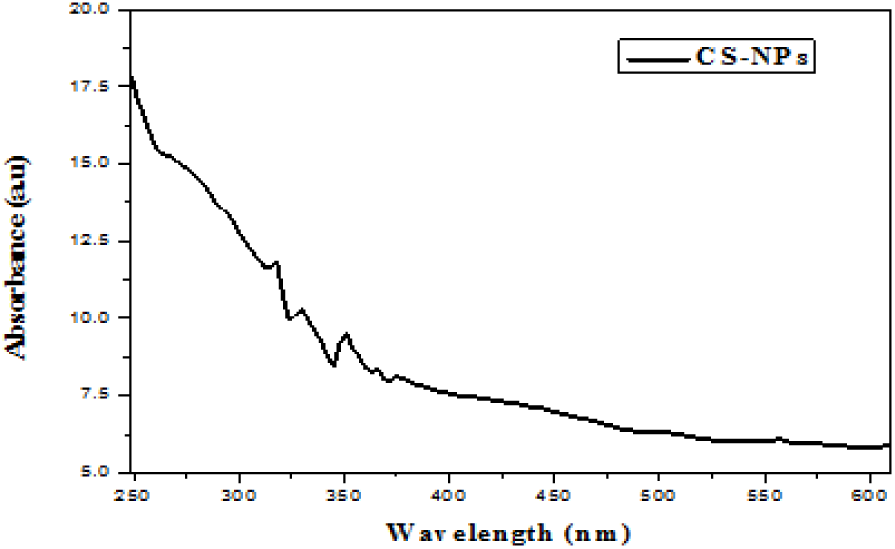
The UV-vis spectra of the biogenic Fe_2_O_3_-CsNPs using *Crocus sativus* corm aqueous extract.

### 2.0.5 Energy Dispersive X-ray Spectroscopic (EDX) Analysis

The scanning electron microscope (JSM5910 JEOL, Japan) was employed for the elemental analysis of biogenic of Fe_2_O_3_-CsNPs. The EDX spectrum revealed the elemental composition of synthesized Fe_2_O_3_-CsNPs Fig 2.0.

**Fig 2.0.**
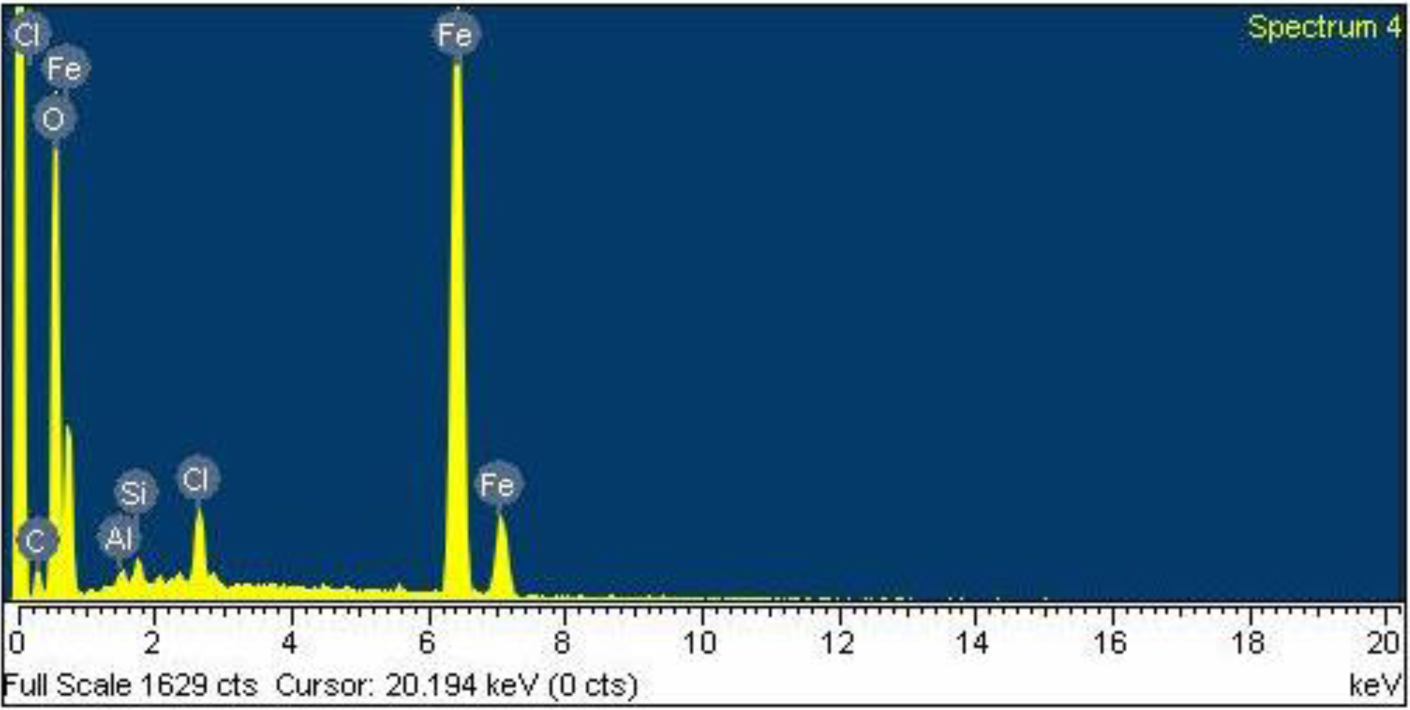
The EXD spectrum of biosynthesized Fe_2_O_3_-CsNPs

### 2.0.6 Scanning Electron Microscopy (SEM) Analysis

The morphology of the acquired Fe_2_O_3_-CsNPs were analyzed, and SEM micrograph were obtained via scanning electron microscope (JSM5910 JEOL, Japan) under different magnifications Fig 3.0.

**Fig. 3.0.**
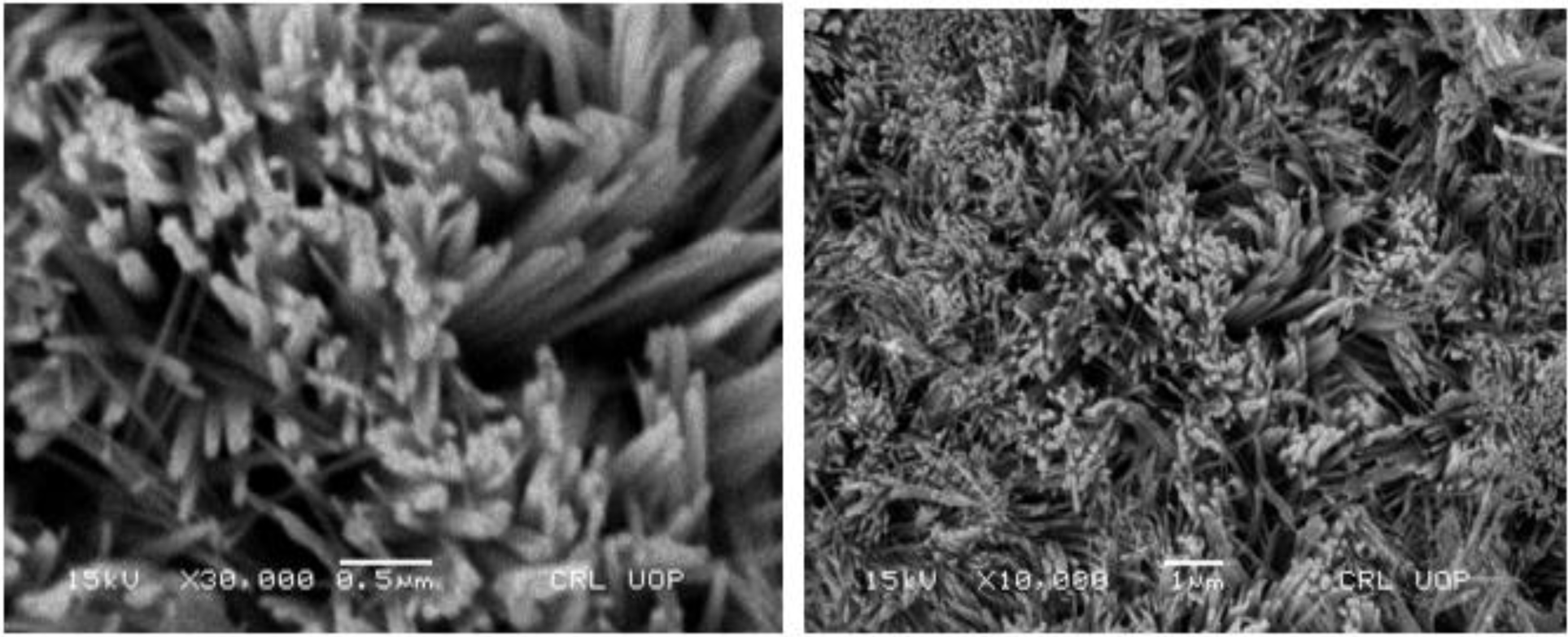
the SEM micrographs of biogenic Fe_2_O_3_-CsNPs

### 2.0.7 Fourier Transform Infrared Spectroscopic (FT-IR) Analysis

The presence of phytochemical and capping of biomolecules on biogenic Fe_2_O_3_-CsNPs surface was analyzed via FT-IR spectrometer (MX BioRad Merlin Ex-Caliber™ FTIR Series, USA). The FT-IR spectrum confirm the presence of numerous biomolecules Fig 4.0.

**Fig 4.0.**
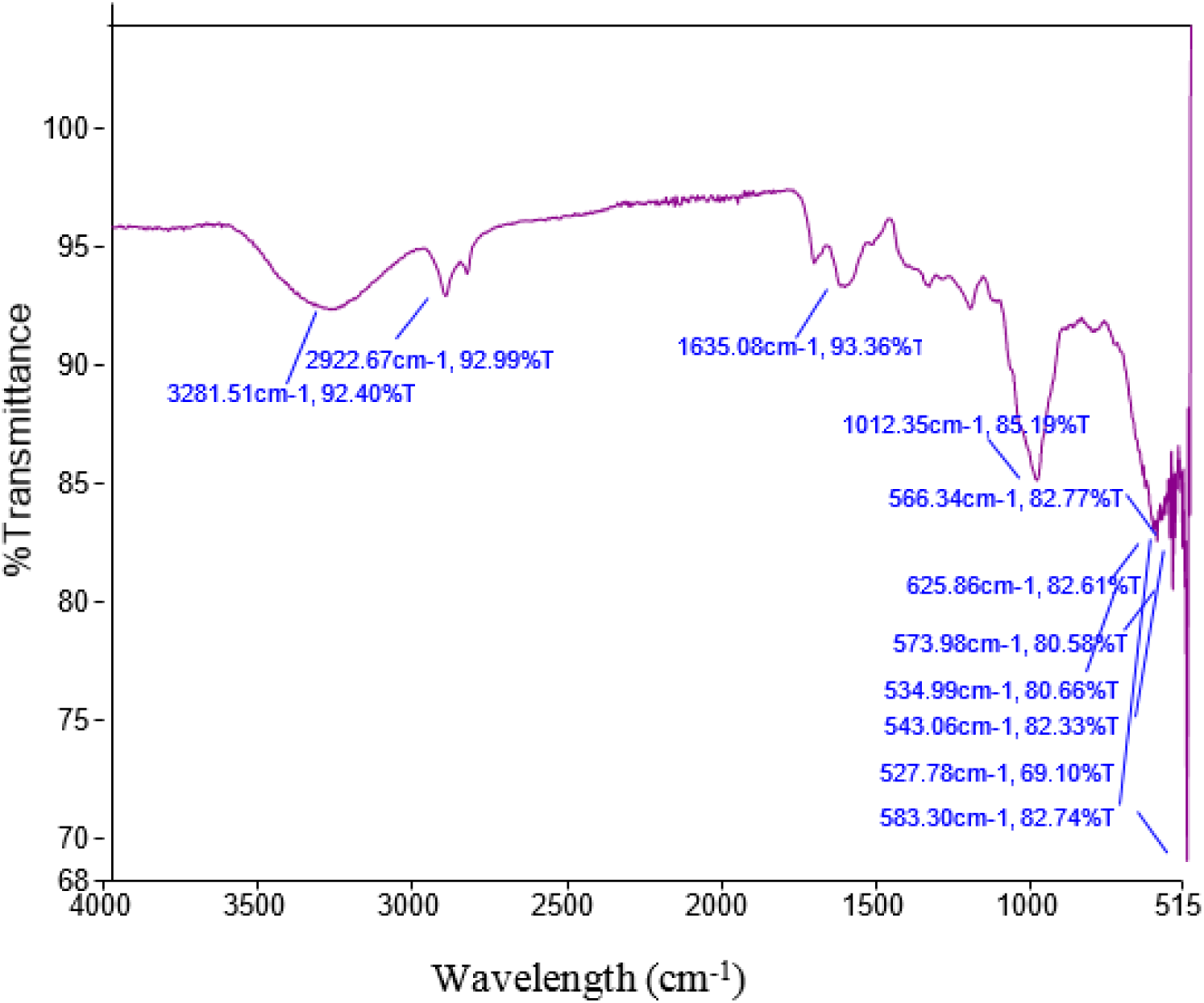
FT-IR spectrum of Fe_2_O_3_-CsNPs

### 2.0.8 X-Ray Diffraction (XRD) Analysis

The size and crystalline texture of the obtained Fe_2_O_3_-CsNPs were analyzed through X-ray diffraction (Siemens D5000 Bruker, Germany) Fig 5.0. The average size of the nanoparticles was computed through the Debye-Scherrer equation.

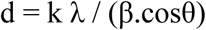

Where *d* is size of Fe_2_O_3_-CsNPs, Sherrer constant is *k*, λ is X-ray wavelength, the width of XRD spectrum at semi height is β, and θ is the Bragg diffraction angle is θ.

**Fig 5.0.**
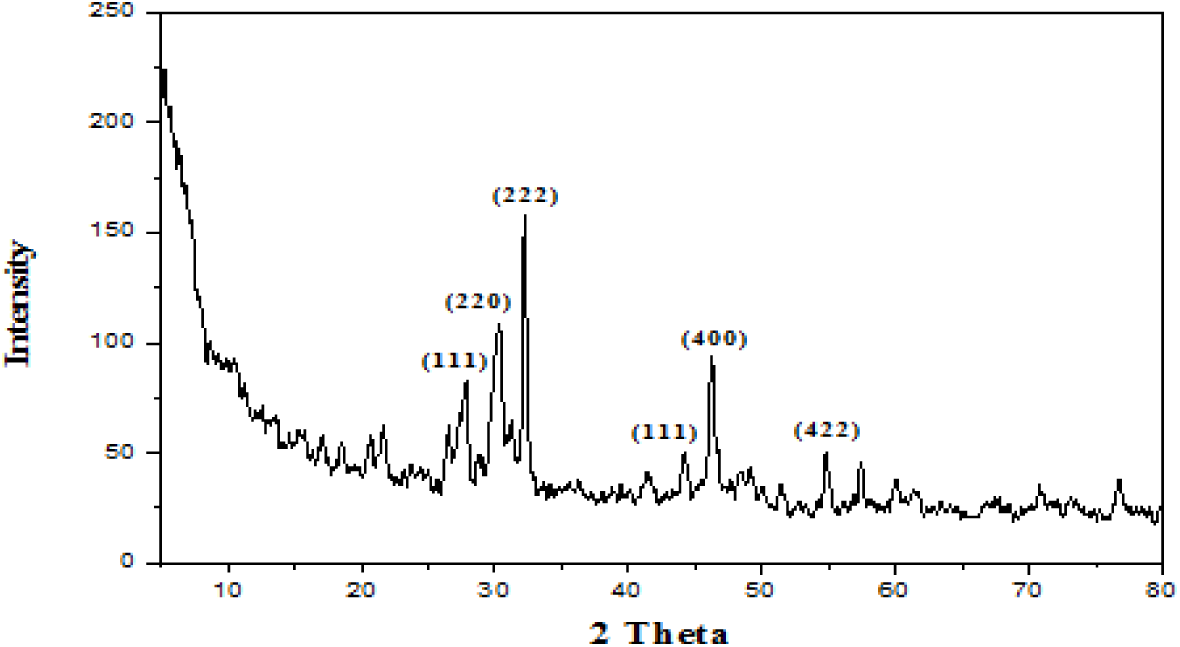
The XRD spectrum of biogenic Fe_2_O_3_-CsNPs via *Crocus sativus* aqueous extract.

### 2.0.9 Antifungal efficacy assay

To evaluate the antifungal efficacy of the nanoparticles on mycelium growth of *V. dahliae*, numerous PDA were prepared including various concentrations of Fe_2_O_3_-CsNPs such as 0.5, 1 and 3 mg/mL [26]. The Petri Plates without the addition of nanoparticles were choosed as positive control. Subsequently, a 6-mm diameter mycelium disk in upside down position was placed in the center of PDA plates and incubated in dark at 25 °C. The mycelium growth was estimated through measuring the colony diameter after 24 h. When the mycelia reached to the margin of positive plates, the antifungal index was computed by the following equation:

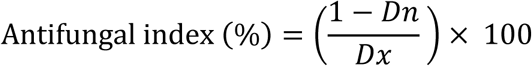

Where *D*_*n*_ is the average mycelia growth diameter in test plate whereas *D*_*x*_ is the average growth diameter of mycelia in control plate [27].

## Results

### 3.0.1 UV-visible Spectroscopic Analysis

The biosynthesis of Fe_2_O_3_-CsNPs was monitored through UV-visible spectroscope. The UV-vis spectra of the synthesized nanoparticles were recorded with 200-800 nm spectral range. The acquired UV-vis spectrum of Fe_2_O_3_-CsNPs is illustrated in Fig 3.0. The characteristic absorption spectra was revealed at 318 nm wavelength which indicated the biogenic reduction of ferric chloride into iron oxide nanoparticles. The results aquired are in accordance with previous reported work [28].

### 3.0.2 X-Ray Diffraction

The average size and morphology of the obtained Fe_2_O_3_-CsNPs were analyzed through XRD as depicted in Fig 5.0. The XRD spectrum showed numerous Bragg’s peaks at 2*θ* =27.76°, 30.24°, 32.19°, 44.26°, 46.19°, 54.80° which can be indexed to 111, 220, 222, 111, 400, 422. The 422 plane indicating the biosynthesis of iron nanoparticles and crystalline morphology (JCPDS Card No.39-1346). The Scherrer’s equation computed average size of the biogenic Fe_2_O_3_-CsNPs was 35-50 nm. The 27.76° peak indicate γ-Fe_2_O_3_ [29]. The results obtained are in good agreement with previous reported work [30].

### 3.0.3 Fourier Transform Infrared Spectroscopic (FT-IR) Analysis

The presence of phytochemical in the Fe_2_O_3_-CsNPs were revealed via FT-IR spectrum. The phytochemical serves as both reducing and stabilizing agents. The acquired FT-IR spectrum depicted various absorption peaks which represent different functional groups Fig 4.0. Various absorption peaks originates at 527, 534, 543, 566, 573, 583, 625, 1012, 1635, 2922 and 3281 cm^− 1^ respectively. The absorption bands less than 800 cm^−1^ are associated with Fe-O and the bending vibration mode of α-Fe_2_O_3_ [31]. The band 1012 cm^−1^ is associated with C-N stretch while the 3281 cm^−1^ is associated with – OH stretch which revealed the reduction of ferric chloride [32]. The band at 2922 cm^−1^ is associated with –C-H stretch while the band at 1635 cm^−1^ is associated with N-H bend [33]. Variation in the absorption peaks were revealed which indicates the oxidation process during the biosynthesis of nanoparticles [34].

### 3.0.4 Energy Dispersive X-ray Spectroscopic Analysis

The EDX configuration revealed the presence of Fe_2_O_3_-CsNPs by depicting Iron and Oxygen spectra Fig 2.0. The hypothesis that the organic moiety play a key role as capping agent was validated with the observation of Carbon spectrum. Cl spectrum was observed which emerged from iron chloride employed to produce Fe_2_O_3_-CsNPs. Identical results has already been obtained under numerous reported work [35, 36]. The weight percentage of Fe, O, C, Cl, Si and Al were 78.32, 15.89, 2.70, 2.18, 0.53, and 0.38 respectively. The presence of Al and Si indicate the impurities. The prominent peaks at 6 and 7 keV showed the presence of elemental iron. The oxygen spectrum indicated the oxidation of nanoparticles due to exposure to air and water [22].

### 3.0.5 Scanning Electron Microscopy (SEM)

The shape and surface morphology of the biogenic Fe_2_O_3_-CsNPs were determined through scanning electron microscopy. The texture and size of Fe_2_O_3_-CsNPs is depicted in SEM micrograph Fig 3.0. The synthesized nanoparticle are crystalline in nature and nanorod in shape. The average size of the acquired Fe_2_O_3_-CsNPs was 58-170 nm. Identical results has already been obtained under numerous reported work [33, 37].

### 3.0.6 Antifungal efficacy assay

The antifungal efficacy of Fe_2_O_3_-CsNPs against *V. dahliae* was analyzed and compared with the *Crocus sativus* corm aqueous extract, and mancozeb at different concentrations (Fig 4.0). The result obtained indicated that the fungus mycelia was significantly degenerated in petri plates contained 0.5, 1 and 3 mg/mL of Fe_2_O_3_-CsNPs with antifungal indexes of 72.1, 77.3, and 86.4% respectively Fig 6.0. Furthermore, the mycelia was greatly reduced as the Fe_2_O_3_-CsNPs concentration enhanced, demonstrating that the antifungal efficacy of Fe_2_O_3_-CsNPs was concentration dependent. The overall results indicates that Fe_2_O_3_-CsNPs exhibited remarkable antifungal effect when compared with *Crocus sativus* corm aqueous extract, and mancozeb under all tested concentrations.

**Fig. 5.**
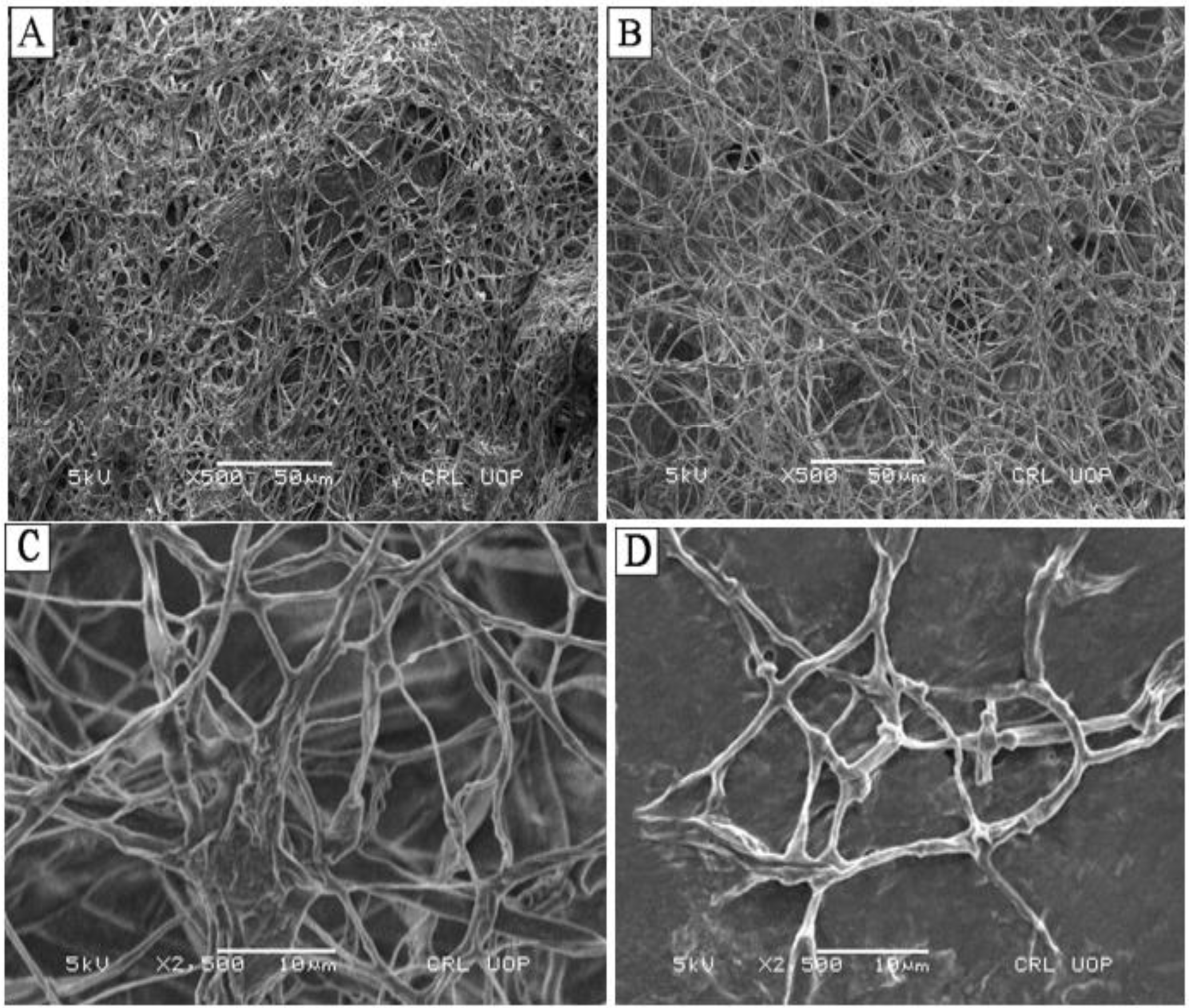
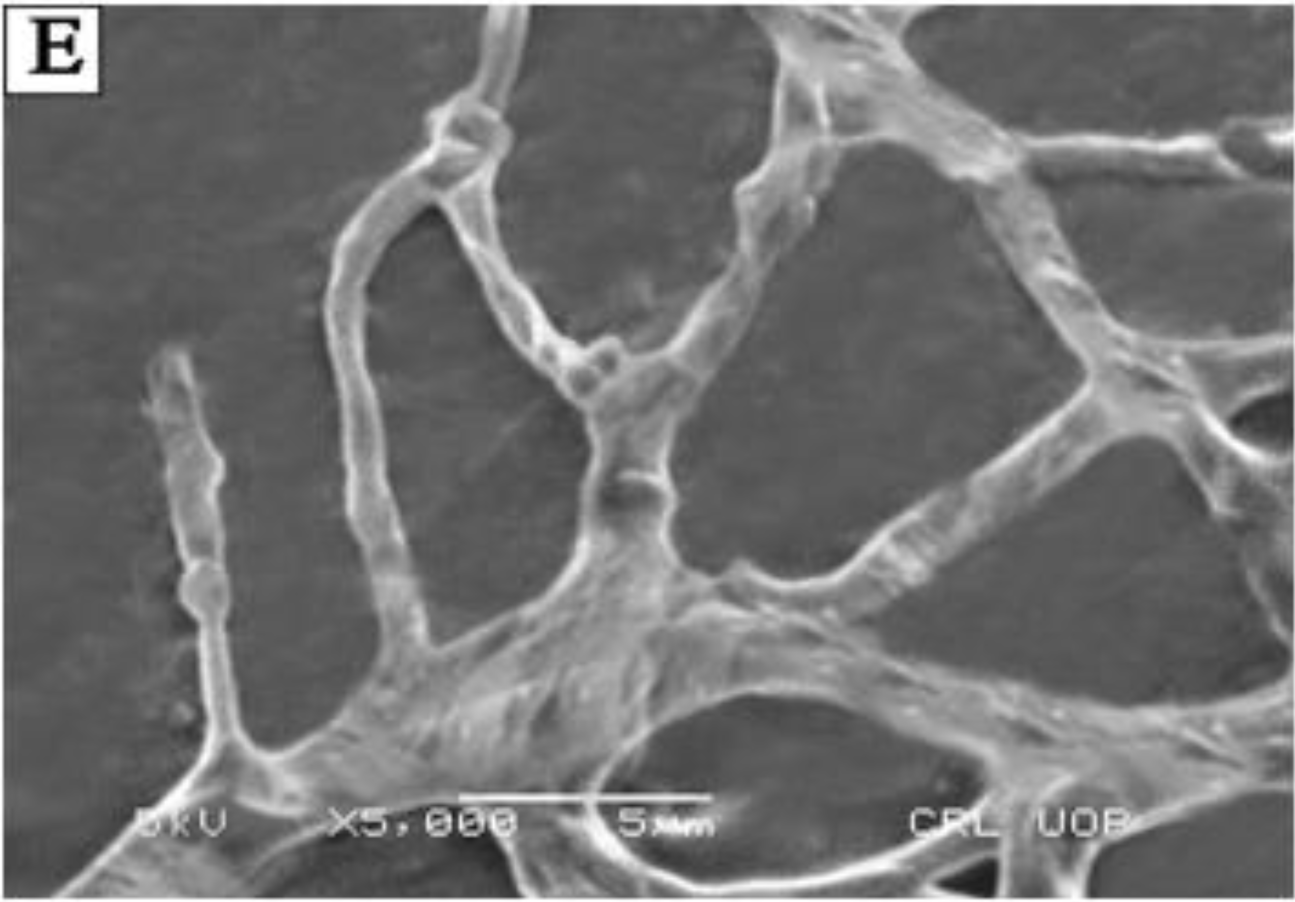
SEM micrographs of *V. dahliae* before the treatment with Fe_2_O_3_-CsNPs (A) and after the treatment with 3 mg/mL of Fe_2_O_3_-CsNPs.

**Fig.6.0.**
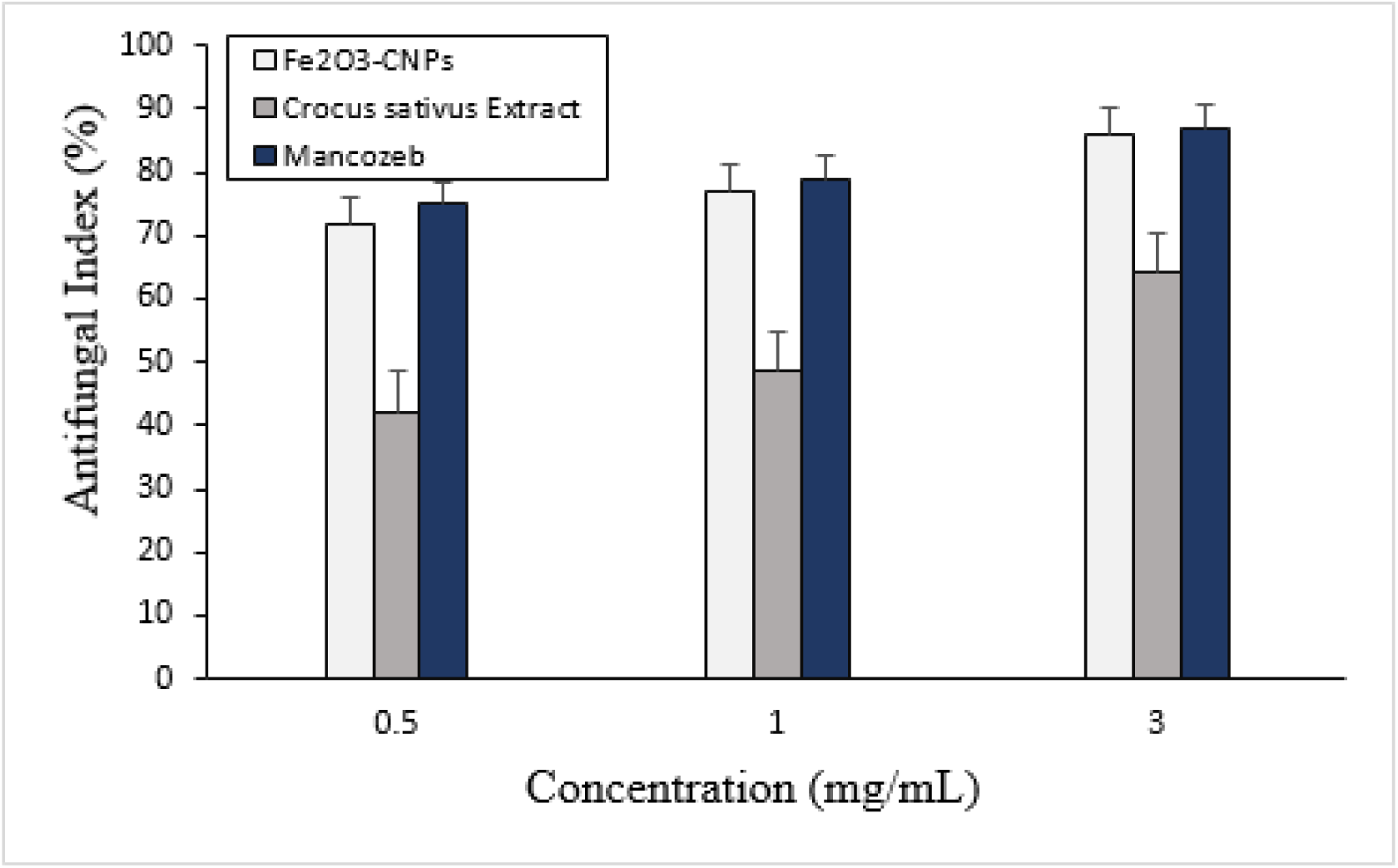
The antifungal index of Fe_2_O_3_-CsNPs, *C. sativus* corm aqueous extract and mancozeb against *V. dahliae* under various concentrations.

## Discussion

In this study, Fe_2_O_3_-CsNPs were employed to evaluate the *in-vitro* antiphytofungal efficacy against *Verticillium dahliae*. The alternation in the mycelium growth of *V. dahliae* indicated that it was sensitive to Fe_2_O_3_-CsNPs in concentration dependent manner, the results obtained were identical to mycelium modification of *V. dahliae* in response to oleoyl-chitosan nanoparticles [38]. Further, Fe_2_O_3_-CsNPs demonstrated advantages over *Crocus sativus* corm aqueous extract under all tested concentrations. The results acquired revealed, Fe_2_O_3_-CsNPs established a clear superiority upon *Crocus sativus* corm extract to inhibit *V. dahliae.*

The SEM micrograph of the biogenic Fe_2_O_3_-CsNPs showed nanorod morphology. The results acquired are identical to previous reported work [33]. The FTIR results are similar to previous reported work on the biochemical constituents in *Crocus sativus* aqueous extract [39]. The UV spectra showed at 318 nm wavelength, similar results have been in previous published work [28]. Likewise, the EDX and XRD results were identical to previously reported work [29, 35, 36].

To investigate the mode of action of Fe_2_O_3_-CsNPs against *V. dahliae*, alternation in hyphae morphology was observed through scanning electron microscopy. The SEM micrograph revealed significant alteration in *V. dahliae* mycelium morphology, the hyphae become degenerated, concentrated and shriveled under the influence of Fe_2_O_3_-CsNPs in treated group Fig. B-E. Nevertheless, the control group showed characteristic morphology i.e., uniform, equal in diameter, smooth and full length hyphae. Cell envelope (i.e., cell membrane and cell wall) shield the leakage of cellular components to extracellular environment. The result of this study demonstrated that the first phase in which Fe_2_O_3_-CsNPs revealed antimicrobial efficacy is due to the interference of nanoparticles with the primary role of cell envelope. With antimicrobial contact, the cytoplasmic membrane may be eliminated or may become functionally inefficient. The destruction of microbial membrane is due to the leakage of micro ions (i.e., potassium and phosphate radical), accompanied by big molecules (i.e., DNA, RNA and other intracellular substance). Consequently, in treatment group acceleration in A260 was observed which imply the direct leakage of cellular membrane and thus a weak membrane [40, 41]. Further, as the Fe_2_O_3_-CsNPs concentration enhanced the total cellular protein concentration reduced. Overall, the results acquired were in good concurrence with morphological observation.

## Conclusion

In this study, Fe_2_O_3_-CsNPs exerted remarkable antiphytofungal efficacy against *V. dahliae*. The antimicrobial effect of Fe_2_O_3_-CsNPs were much stronger than *C. sativus* corm aqueous extract in all tested concentrations. Fe_2_O_3_-CsNPs significantly prevent the proliferation of mycelium. The incorporation of nanoparticles engendered to release of intracellular substances, consequently lead to the depressive alternation of external texture. These results demonstrated that Fe_2_O_3_-CsNPs have the efficacy to control devastative phytopathogens. Nevertheless, the mechanism of antimicrobial efficacy might be more perplexed than the finding of this study appears to suggest. Further studies should investigate the antimicrobial efficacy of Fe_2_O_3_-CsNPs for the control of numerous plant diseases.

## References

1. Atallah, Z.K., R.J. Hayes, and K.V. Subbarao, Fifteen years of Verticillium wilt of lettuce in America’s salad bowl: a tale of immigration, subjugation, and abatement. Plant disease, 2011. 95(7): p. 784–792.

2. Bhat, R. and K. Subbarao, Host range specificity in Verticillium dahliae. Phytopathology, 1999. 89(12): p. 1218–1225.

3. Lu, W., et al., Verticillium wilt of redbud in China caused by Verticillium dahliae. Plant Disease, 2013. 97(11): p. 1543.

4. Sun, B., et al., Agricultural non-point source pollution in China: causes and mitigation measures. Ambio, 2012. 41(4): p. 370–379.

5. Park, H.-J., et al., A new composition of nanosized silica-silver for control of various plant diseases. The plant pathology journal, 2006. 22(3): p. 295–302.

6. Raghupathi, K.R., R.T. Koodali, and A.C. Manna, Size-dependent bacterial growth inhibition and mechanism of antibacterial activity of zinc oxide nanoparticles. Langmuir, 2011. 27(7): p. 4020–4028.

7. Kim, S.W., et al., Antifungal effects of silver nanoparticles (AgNPs) against various plant pathogenic fungi. Mycobiology, 2012. 40(1): p. 53–58.

8. Alam, T., et al., Biogenic synthesis of iron oxide nanoparticles via Skimmia laureola and their antibacterial efficacy against bacterial wilt pathogen Ralstonia solanacearum. 2019. 98: p. 101–108.

9. Parsons, J., J. Peralta-Videa, and J. Gardea-Torresdey, Use of plants in biotechnology: synthesis of metal nanoparticles by inactivated plant tissues, plant extracts, and living plants. Developments in environmental science, 2007. 5: p. 463–485.

10. Shameli, K., et al., Green biosynthesis of silver nanoparticles using Curcuma longa tuber powder. International journal of nanomedicine, 2012. 7: p. 5603.

11. Jain, T.K., et al., Iron oxide nanoparticles for sustained delivery of anticancer agents. Molecular pharmaceutics, 2005. 2(3): p. 194–205.

12. Wang, Z., C. Fang, and M. Megharaj, Characterization of iron–polyphenol nanoparticles synthesized by three plant extracts and their fenton oxidation of azo dye. ACS Sustainable Chemistry & Engineering, 2014. 2(4): p. 1022–1025.

13. Wang, T., et al., Green synthesis of Fe nanoparticles using eucalyptus leaf extracts for treatment of eutrophic wastewater. Science of the total environment, 2014. 466: p. 210–213.

14. Mohanpuria, P., N.K. Rana, and S.K. Yadav, Biosynthesis of nanoparticles: technological concepts and future applications. Journal of nanoparticle research, 2008. 10(3): p. 507–517.

15. Seabra, A.B., P. Haddad, and N. Duran, Biogenic synthesis of nanostructured iron compounds: applications and perspectives. IET nanobiotechnology, 2013. 7(3): p. 90–99.

16. Barnes, R.J., et al., The impact of zero-valent iron nanoparticles on a river water bacterial community. Journal of hazardous materials, 2010. 184(1-3): p. 73–80.

17. El-Temsah, Y.S. and E.J. Joner, Impact of Fe and Ag nanoparticles on seed germination and differences in bioavailability during exposure in aqueous suspension and soil. Environmental toxicology, 2012. 27(1): p. 42–49.

18. Kumar, K.M., et al., Biobased green method to synthesise palladium and iron nanoparticles using Terminalia chebula aqueous extract. Spectrochimica Acta Part A: Molecular and Biomolecular Spectroscopy, 2013. 102: p. 128–133.

19. Madhavi, V., et al., Application of phytogenic zerovalent iron nanoparticles in the adsorption of hexavalent chromium. Spectrochimica Acta Part A: Molecular and Biomolecular Spectroscopy, 2013. 116: p. 17–25.

20. López-Téllez, G., et al., Green method to form iron oxide nanorods in orange peels for chromium (VI) reduction. Journal of nanoscience and nanotechnology, 2013. 13(3): p. 2354–2361.

21. Chrysochoou, M., C.P. Johnston, and G. Dahal, A comparative evaluation of hexavalent chromium treatment in contaminated soil by calcium polysulfide and green-tea nanoscale zero-valent iron. Journal of hazardous materials, 2012. 201: p. 33–42.

22. Njagi, E.C., et al., Biosynthesis of iron and silver nanoparticles at room temperature using aqueous sorghum bran extracts. Langmuir, 2010. 27(1): p. 264–271.

23. Serrano-Diaz, J., et al., Flavonoid determination in the quality control of floral bioresidues from Crocus sativus L. Journal of agricultural and food chemistry, 2014. 62(14): p. 3125–3133.

24. Khazaei, K.M., et al., Application of maltodextrin and gum Arabic in microencapsulation of saffron petal’s anthocyanins and evaluating their storage stability and color. Carbohydrate polymers, 2014. 105: p. 57–62.

25. Goupy, P., et al., Identification and quantification of flavonols, anthocyanins and lutein diesters in tepals of Crocus sativus by ultra performance liquid chromatography coupled to diode array and ion trap mass spectrometry detections. Industrial crops and products, 2013. 44: p. 496–510.

26. Palma-Guerrero, J., et al., Effect of chitosan on hyphal growth and spore germination of plant pathogenic and biocontrol fungi. Journal of applied Microbiology, 2008. 104(2): p. 541–553.

27. Guo, Z., et al., Novel derivatives of chitosan and their antifungal activities in vitro. Carbohydrate Research, 2006. 341(3): p. 351–354.

28. Devatha, C., A.K. Thalla, and S.Y. Katte, Green synthesis of iron nanoparticles using different leaf extracts for treatment of domestic waste water. Journal of cleaner production, 2016. 139: p. 1425–1435.

29. Weng, X., et al., Synthesis of iron-based nanoparticles by green tea extract and their degradation of malachite. Industrial Crops and Products, 2013. 51: p. 342–347.

30. Patra, J.K. and K.-H. Baek, Green biosynthesis of magnetic iron oxide (Fe3O4) nanoparticles using the aqueous extracts of food processing wastes under photo-catalyzed condition and investigation of their antimicrobial and antioxidant activity. Journal of Photochemistry and Photobiology B: Biology, 2017. 173: p. 291–300.

31. Sahoo, S., et al., Characterization of γ-and α-Fe 2 O 3 nano powders synthesized by emulsion precipitation-calcination route and rheological behaviour of α-Fe 2 O 3. International Journal of Engineering, Science and Technology, 2010. 2(8).

32. Litvin, V.A., B.F. Minaev, and G.V. Baryshnikov, Synthesis and properties of synthetic fulvic acid derived from hematoxylin. Journal of Molecular Structure, 2015. 1086: p. 25–33.

33. Rajiv, P., et al., Synthesis and characterization of biogenic iron oxide nanoparticles using green chemistry approach and evaluating their biological activities. Biocatalysis and Agricultural Biotechnology, 2017. 12: p. 45–49.

34. Vidhu, V. and D. Philip, Spectroscopic, microscopic and catalytic properties of silver nanoparticles synthesized using Saraca indica flower. Spectrochimica Acta Part A: Molecular and Biomolecular Spectroscopy, 2014. 117: p. 102–108.

35. Machado, S., et al., Characterization of green zero-valent iron nanoparticles produced with tree leaf extracts. Science of the total environment, 2015. 533: p. 76–81.

36. Wang, T., et al., Green synthesized iron nanoparticles by green tea and eucalyptus leaves extracts used for removal of nitrate in aqueous solution. Journal of cleaner production, 2014. 83: p. 413–419.

37. Mahdavi, M., et al., Green biosynthesis and characterization of magnetic iron oxide (Fe3O4) nanoparticles using seaweed (Sargassum muticum) aqueous extract. Molecules, 2013. 18(5): p. 5954–5964.

38. Xing, K., et al., Effect of O-chitosan nanoparticles on the development and membrane permeability of Verticillium dahliae. Carbohydrate polymers, 2017. 165: p. 334–343.

39. Bagherzade, G., M.M. Tavakoli, and M.H. Namaei, Green synthesis of silver nanoparticles using aqueous extract of saffron (Crocus sativus L.) wastages and its antibacterial activity against six bacteria. Asian Pacific Journal of Tropical Biomedicine, 2017. 7(3): p. 227–233.

40. Chen, C.Z. and S.L. Cooper, Interactions between dendrimer biocides and bacterial membranes. Biomaterials, 2002. 23(16): p. 3359–3368.

41. Xing, K., et al., Antibacterial activity of oleoyl-chitosan nanoparticles: A novel antibacterial dispersion system. Carbohydrate polymers, 2008. 74(1): p. 114–120.

